# Intratumoral immune triads are required for adoptive T cell therapy-mediated elimination of solid tumors

**DOI:** 10.1101/2023.07.03.547423

**Authors:** Gabriel Espinosa-Carrasco, Aurora Scrivo, Paul Zumbo, Asim Dave, Doron Betel, Matthew Hellmann, Bryan M Burt, Hyun-Sung Lee, Andrea Schietinger

## Abstract

Tumor-reactive CD8 T cells found in cancer patients are frequently dysfunctional, unable to halt tumor growth. Adoptive T cell transfer (ACT), the administration of large numbers of *in vitro*-generated cytolytic tumor-reactive CD8 T cells, is an important cancer immune therapy being pursued. However, a limitation of ACT is that transferred CD8 T cells often rapidly lose effector function, and despite exciting results in certain malignancies, few ACT clinical trials have shown responses in solid tumors. Here, we developed preclinical cancer mouse models to investigate if and how tumor-specific CD4 T cells can be enlisted to overcome CD8 T cell dysfunction in the setting of ACT. *In situ* confocal microscopy of color-coded cancer cells, tumor-specific CD8 and CD4 T cells, and antigen presenting cells (APC), combined with functional studies, revealed that the spatial positioning and interactions of CD8 and CD4 T cells, but not their numbers, dictates ACT efficacy and anti-tumor responses. We uncover a new role of antigen-specific CD4 T cells in addition to the known requirement for CD4 T cells during priming/activation of naïve CD8 T cells. CD4 T cells must co-engage with CD8 T cells and APC cross-presenting CD8-and CD4-tumor antigens during the effector phase, forming a three-cell-cluster (triad), to license CD8 T cell cytotoxicity and mediate cancer cell elimination. Triad formation transcriptionally and epigenetically reprogram CD8 T cells, prevent T cell dysfunction/exhaustion, and ultimately lead to the elimination of large established tumors and confer long-term protection from recurrence. When intratumoral triad formation was disrupted, adoptively transferred CD8 T cells could not be reprogrammed, and tumors progressed despite equal numbers of tumor-infiltrating CD8 and CD4 T cells. Strikingly, the formation of CD4 T cell::CD8 T cell::APC triads in tumors of patients with lung cancers treated with immune checkpoint blockade was associated with clinical responses, but not CD4::APC dyads or overall numbers of CD8 or CD4 T cells, demonstrating the importance of triads in non-ACT settings in humans. Our work uncovers intratumoral triads as a key requirement for anti-tumor immunity and a new role for CD4 T cells in CD8 T cell cytotoxicity and cancer cell eradication.

## INTRODUCTION

CD8 T cells are powerful components of the adaptive immune system that have the potential to selectively eradicate cancer cells. However, despite the presence of tumor-specific CD8 T cells in tumor-bearing hosts, cancers develop, suggesting that CD8 T cells become dysfunctional and unresponsive to cancer cells over the course of tumorigenesis [1]. Tumor-infiltrating dysfunctional CD8 T cells (also referred to as ‘exhausted’ T cells) commonly express high levels of inhibitory receptors (PD1, LAG3, CTLA4, TIM3) and fail to produce effector cytokines (interferon-γ (IFN-γ), tumor necrosis factor-α (TNF-α)) and cytotoxic molecules (granzymes, perforin). These hallmarks of CD8 T cell dysfunction/exhaustion have been attributed to chronic tumor antigen encounter/TCR signaling and immunosuppressive signals within the tumor microenvironment [1-3].

Adoptive T cell transfer (ACT), the infusion of large numbers (> 10^9^ – 10^10^ CD8 T cells/patient) of tumor-reactive cytolytic effector CD8 T cells into cancer patients, has emerged as a powerful therapeutic strategy for the treatment of cancers [4]. Tumor-reactive CD8 T cells can either be isolated from patients’ own tumors (tumor-infiltrating lymphocytes (TIL)) or blood, expanded *ex vivo* and infused back, or engineered *in vitro* to become tumor-reactive through the introduction of genes encoding T cell receptors (TCR) or chimeric antigen receptors (CAR) specific for tumor antigens [5-11]. Although remarkable successes with ACT have been observed in a subset of cancer patients and cancer types (e.g. leukemia, lymphoma, and melanoma) [12-14], most patients still fail to achieve long-term responses, especially those with (non-melanoma) solid tumors. Factors which mitigate the efficacy of adoptively transferred CD8 T cells include poor *in vivo* persistence, poor tumor localization/infiltration, and rapid loss of effector function [13, 15, 16]. Various therapeutic strategies have been identified to improve persistence and tumor infiltration, such as lymphodepletion and/or administration of homeostatic cytokines (IL-2, IL-7, IL-15) [12, 15, 17-21]. However, the loss of effector function of CD8 T cells remains a major roadblock [22, 23]. Thus, the development of immunotherapeutic interventions to prevent or reverse CD8 T cell dysfunction/exhaustion has become the concerted effort of many clinicians and scientists.

While direct cytotoxic activity against cancer cells generally resides within the CD8 T cell compartment, various modes of action have been described for CD4 T cells [24]: (1) productive priming of naïve CD8 T cells in lymphoid tissues through “licensing” and functional maturation of dendritic cells (DC) [25-31], (2) anti-tumor effector functions and elimination of MHC class II-negative cancer cells without CD8 T cells [32-36] through IFN-γ acting on the host stroma, or activation of macrophages and other non-lymphoid tumoricidal effector cells [35, 37-42], and (3) induction of cancer cell senescence rather than cancer cell elimination through the secretion of Th1-cytokines (TNFα, IFNγ) [43, 44]. Moreover, we and others have demonstrated that CD4 T cells might play an important role during CD8 T cell-mediated tumor elimination as well as during autoimmune tissue destruction, however, the mechanisms remained elusive [45-47]. MHC class II-restricted tumor antigens and tumor-specific CD4 T cells have been identified in many cancer patients and cancer types, and their importance in anti-tumor immunity has been recognized [24, 32, 48-52]. If and how tumor-reactive CD4 T cells can be utilized to prevent or reverse CD8 T cell dysfunction/exhaustion leading to tumor eradication is not known. To address this question, we developed a clinically relevant ACT-cancer mouse model. We demonstrate that CD4 T cells mediate tumor-specific CD8 T cell reprogramming within large solid tumors when tumor-reactive CD4 and CD8 T cells form three-cell-type clusters (triads) together with antigen-presenting cells (APC). Triad-formation resulted in the molecular and functional reprogramming of adoptively transferred CD8 T cells, preventing and even reversing T cell exhaustion, leading to tumor destruction. Strikingly, the formation of CD4 T cell-CD8 T cell-APC triads in tumors of patients with mesothelioma treated with immune checkpoint blockade (ICB) was associated with clinical responses, uncovering CD4 T cell-CD8 T cell-APC triads as a key determinant for cancer elimination and ACT therapy efficacy against solid tumors.

## RESULTS

### Tumor-specific CD4 T cells reverse tumor-specific CD8 T cell dysfunction/exhaustion in solid tumors

B16 is a highly aggressive murine melanoma cell line; B16 cancer cells injected subcutaneously into immunocompetent C57BL/6 wildtype mice (B6 WT) form large established tumors within 2 weeks, ultimately killing the host, and treatment regiments are generally ineffective. We engineered B16 cancer cells to express the CD8 T cell–recognized epitope from ovalbumin OVA_257-264_ (SIINFEKL) as well as the CD4 T cell-recognized glycoprotein epitope GP_61–80_ (GLKGPDIYKGVYQFKSVEFD) from the lymphocytic choriomeningitis virus (LCMV); the vector was constructed to encode the trimeric peptide sequence (SIINFEKL-AAY)_3_ fused to the fluorescent protein Cerulean, followed by the 19-mer GP_61–80_ peptide (**Fig.1a**). The OVA_257-264_ epitope is presented on the MHC class I molecule H-2K^b^ and recognized by TCR transgenic OT1 CD8 T cells (TCR_OT1_); the GP_61–80_ epitope is presented on the MHC class II I-A^b^ molecule and recognized by TCR transgenic SMARTA CD4 T cells (TCR_SMARTA_). B16-OVA_257-264_-GP_61– 80_ cancer cells (B16-OG; 2.5 x10^6^ cells/host) were injected subcutaneously into B6 WT (CD45.2) mice. Despite the expression of strong CD8-and CD4-T cell tumor antigens, B16-OG tumors grew aggressively, forming large tumors within 2 weeks (**Fig. 1b**). We then employed an adoptive T cell transfer (ACT) regimen modeled on that used in cancer patients treated with ACT: preconditioning the host and inducing lymphopenia through a nonmyeloablative chemotherapeutic dose of cyclophosphamide followed by the infusion of *in vitro* generated cytotoxic tumor-specific CD8 T cells (**Fig. 1a**). Naïve congenic (CD45.1) TCR_OT1_ were activated *in vitro* for 3-4 days and adoptively transferred into lymphopenic B16-OG tumor-bearing mice. Despite the infusion of highly functional effector TCR_OT1_ CD8 T cells, B16-OG tumors progressed, recapitulating the scenario commonly observed in patients with solid tumors receiving ACT (**Fig. 1b**). Next, we asked whether the simultaneous infusion of *in vitro* activated effector TCR_SMARTA_ CD4 T cells would mediate anti-tumor responses. Co-transfer of effector TCR_OT1_ together with TCR_SMARTA_ resulted in complete tumor elimination, with 100% long-term tumor-free survival (**Fig. 1b**). Tumor-bearing mice that received TCR_SMARTA_ alone did not show tumor regression (data not shown), demonstrating that cancer elimination was dependent on both TCR_OT1_ and TCR_SMARTA_ T cells. We confirmed our results in a second tumor model using the fibrosarcoma cell line MCA205 (MCA205-OG) and obtained similar results (**Fig. 1c**).

**Figure 1.**
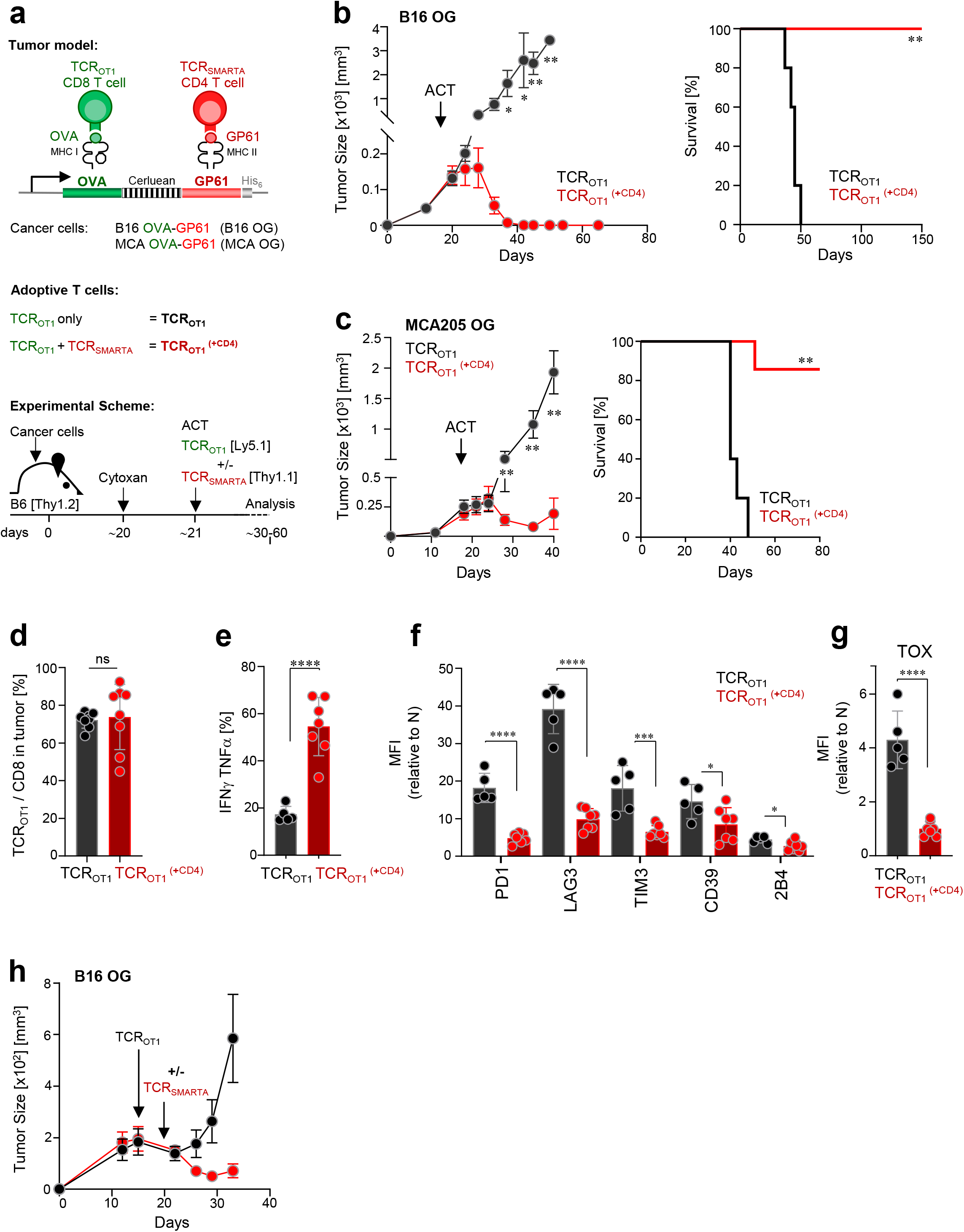
Tumor-specific CD4 T cells prevent and reverse CD8 T cell dysfunction/exhaustion within solid tumors and mediate tumor elimination. **a.** Scheme: tumor models, adoptively transferred effector T cells, and experimental schemes. **b.** B16 OVA-GP_61-80_ (B16-OG) tumor growth (right) and Kaplan–Meier survival curve (left) of tumor-bearing B6 WT mice (CD45.2; Thy1.2) receiving effector TCR_OTI_ CD8 T cells alone (CD45.1) (black; TCR_OT1_) or together with TCR_SMARTA_ CD4 T cells (Thy1.1) (red; TCR ^(+CD4)^) (ACT = adoptive T cell transfer). Data is representative of 5 independent experiments (n=5 mice/group). Values are mean ± SEM. Significance is calculated by multiple *t* test. Kaplan–Meier curve; **p=0.00021; Mantel–Cox test. **c.** MCA205 OVA-GP_61-80_ (MCA-OG) tumor outgrowth and survival in B6 mice treated as described in b; **p=0.0003; Mantel–Cox test. Data is representative of 2 independent experiments (n=5-6 mice/group). **d.** TCR_OTI_ (% of total of CD8+ T cells) within progressing B16-OG tumors 8-9 days post transfer +/- TCR_SMARTA_ CD4 T cells. Data pooled from 2 independent experiments (n=8 mice/group). Each symbol represents an individual mouse. **e.** IFNγ and TNFα production of TCR_OTI_ isolated from B16-OG tumors 8-9 days post transfer +/- TCR_SMARTA_ CD4 T cells. Cytokine production was assessed after 4-hr peptide stimulation *ex vivo*. Data show 2 pooled independent experiments (n=5-7). **f.** Inhibitory receptor expression, and **g.** TOX expression of B16-OG tumor-infiltrating TCR_OTI_ isolated 8-9 days post transfer +/- TCR_SMARTA_. Graphs depict relative MFI normalized to naive TCR_OTI_; two pooled independent experiments (n=5-7mice/group). **h.** Mice with B16-OG tumors received effector TCR_OTI_ CD8 T cells 14 days post tumor transplantation; 9 days later, TCR_SMARTA_ CD4 T cells were adoptively transferred (red); B16-OG tumor growth in mice receiving only TCR_OT1_ are shown in black. Data is representative of 2 independent experiments (n=8 mice/group). Values are mean ± SEM. Significance is calculated by multiple *t* test.

CD4 T cells are known to enhance CD8 T cell mobilization into peripheral tissues [28]. To understand whether TCR_SMARTA_ enhanced TCR_OT1_ tumor infiltration, we compared the numbers of TCR_OT1_ TIL in mice which received effector TCR_OT1_ alone (TCR_OT1_) or together with TCR_SMARTA_ (TCR ^(+CD4)^); we evaluated numbers of TIL 8-9 days post transfer, a time point when tumors are similar in size. Surprisingly, we found equal numbers of TCR_OT1_ TIL in both cohorts (**Fig. 1d**), suggesting that TCR_SMARTA_-mediated anti-tumor immunity was not due to an enhancement of TCR_OT1_ tumor infiltration but likely due to functional changes of TCR_OT1_ TIL. Indeed, while TCR_OT1_ TIL were impaired in their ability to produce the effector cytokines IFNγ and TNFα (**Fig. 1e**), expressed high levels of numerous canonical inhibitory receptors including PD1, LAG3, TIM3, CD39 and 2B4 (**Fig. 1f**), as well as the transcription factor TOX (**Fig. 1g**), a critical regulator associated with T cell exhaustion [53-58], TCR ^(+CD4)^ were able to produce high amounts of IFNγ and TNFα and showed little/no expression of inhibitory receptors and TOX (**Fig. 1e-1g**). To understand whether these phenotypic and functional differences were already induced in the tumor-draining lymph node (tdLN), we compared phenotype and function of tdLN-TCR_OT1_ and tdLN-TCR_OT1_^(+CD4)^. Interestingly, no differences were observed (**Suppl. Fig. 1**), thus co-transferred CD4 T cells specifically acted on tumor-specific CD8 T cells within the tumor.

Next, we wanted to understand whether CD4 T cells could not only prevent but also reverse CD8 T cell dysfunction/exhaustion. We adoptively transferred effector TCR_OT1_ into B16-OG tumor-bearing mice, and 10 days later, when TCR_OT1_ TIL were dysfunctional/exhausted, we adoptively transferred effector TCR_SMARTA_. Remarkably, mice that received TCR_SMARTA_ showed tumor regression while control cohorts did not (**Fig. 1h**). Thus, tumor-reactive TCR_SMARTA_ CD4 T cells prevent and reverse tumor-induced CD8 T cell dysfunction and mediate tumor regression.

### CD4 T cells transcriptionally and epigenetically reprogram tumor-specific CD8 T cells, leading to tumor elimination

Tumor-specific CD8 T cell dysfunction in mice and humans is associated with global transcriptional and epigenetic dysregulation of genes and pathways important for T cell differentiation and function. To understand how CD4 T cells mediated functional rescue of TCR_OT1_ CD8 T cells, we conducted RNA-seq and ATAC-seq of TCR_OT1_^(+CD4)^ and TCR TIL isolated from size-matched B16-OG tumors 8 days post transfer. 1795 genes were differentially expressed (DEG) including exhaustion/dysfunction-associated TF and inhibitory receptors/activation markers (*Tox*, *Irf4*, *Pdcd1* (PD1), *Havcr2*, *Lag3*, *CD160*, *Cd244* (2B4)) (**Fig. 2a** and **2b**), which were highly expressed in TCR_OT1_. In contrast, TF and molecules associated with stem-like progenitor T cell states were enriched and highly expressed in TCR_OT1_^(+CD4)^ TIL, including genes encoding *Tcf7* (TCF1), *Il7r*, *Itgae* (CD103), *Itga1*, and *Ifitm3*, as well as chemokine receptors such as *Ccr5*, *Ccr4* and *Ccr2* [30, 59]. Gene ontology (GO) classification revealed that pathways associated with positive cytokine regulation, immune differentiation and responses to tumor cells were enriched in TCR_OT1_^(+CD4)^ but not in TCR_OT1_ (**Fig. 2c**). ATAC-seq revealed 11,787 differentially accessible regions (DAR), including enhancers in many exhaustion (*Tox*, *Spry1 Spry2, Cd244, Bach2, Egr2*) or stem-/progenitor cell state-associated genes (*Tcf7*, *IL7r*, *Lef1*), respectively (**Fig. 2d** and **2e**). Many enhancer peaks with TF motifs associated with terminal differentiation were less accessible in reprogrammed CD8 T cells, which was surprising given that TCR_OT1_^(+CD4)^ and TCR TIL were isolated from equally sized tumors (**Fig. 2f**). To understand whether reprogrammed TCR_OT1_^(+CD4)^ revealed molecular signatures similar to human CD8 TIL driving clinical responses in the context of ACT, we utilized a data set from a study conducted by the Rosenberg group, using *ex vivo*-expanded autologous CD8+ TIL from metastatic melanoma lesions for ACT into preconditioned, lymphodepleted patients [60]. The authors identified a CD39-CD69-stem-like TIL subset that was associated with complete cancer regression in ACT-responders but lacking in ACT-non-responders. Gene set enrichment analysis (GSEA) revealed that the same genes were enriched in TCR_OT1_^(+CD4)^ CD8 TIL as in ACT (CD39-CD69-) CD8 TIL responders, and genes in CD8 TIL from ACT (CD39+CD69+) non-responders were enriched in TCR_OT1_ CD8 TIL (**Fig. 2g**, **Suppl. Fig. 2**) [60].

Taken together, tumor-specific TCR_SMARTA_ CD4 T cells transcriptionally and epigenetically reprogram tumor-reactive CD8 TIL within progressing tumors, preventing terminal differentiation and exhaustion, and resulting in tumor elimination.

### Spatial positioning of tumor-specific CD8 and CD4 T cells within tumors determine anti-tumor immunity and cancer elimination

Next, we wanted to understand *how* TCR_SMARTA_ CD4 T cells prevent CD8 T cell exhaustion within tumors. B16 tumor cells express low level MHC II *in vivo* (**Suppl. Fig. 3a**), thus cancer cells could become targets of CD4 T cells. Employing CRISPR/Cas9-mediated gene editing, we generated MHC class II *I-A^b^*-deficient B16-OG cancer cells. Surprisingly, large established B16-OG *I-A^b^*-deficient tumors were eliminated as efficiently as parental MHC class II-expressing B16-OG tumors, demonstrating that cancer elimination does not require CD4 T cell to directly target cancer cells (**Suppl. Fig. 3b** and **3c**). Next, we turned to the tumor stroma, which includes MHC class I-and II-expressing antigen presenting cells (APC) such as CD11c+ dendritic cells (DC) and macrophages. To assess the role of CD11c+ cells, we employed a targeted depletion approach: CD11c+ DC from CD11c-DTR/GFP transgenic mice express the primate diphtheria toxin receptor (DTR) transgene under the CD11c promoter, enabling conditional depletion of CD11c+ cells *in vivo* upon DT treatment [61]. We generated bone marrow (BM) chimeras by transferring BM cells from CD11c-DTR/GFP (CD11c-DTR) or littermate control (WT) mice into lethally irradiated WT (CD45.1) B6 mice (designated “DTR→WT” and “WT→WT” chimeras). B16-OG tumors were established in DTR→WT and WT→WT BM chimeras, and 2-3 weeks post B16-OG tumor cell transplantation effector TCR_OTI_ and TCR_SMARTA_ were adoptively transferred. 5 days post ACT, when TCR_OTI_ and TCR_SMARTA_ infiltrated into tumors, mice were treated twice weekly with DT. Depletion of CD11c+ APC prevented tumor elimination in DTR→WT mice but not control WT→WT mice, suggesting that CD11c+ APC within the tumor microenvironment were necessary for TCR_SMARTA_-mediated TCR_OTI_ reprogramming and tumor elimination (**Fig. 3a**).

**Figure 2.**
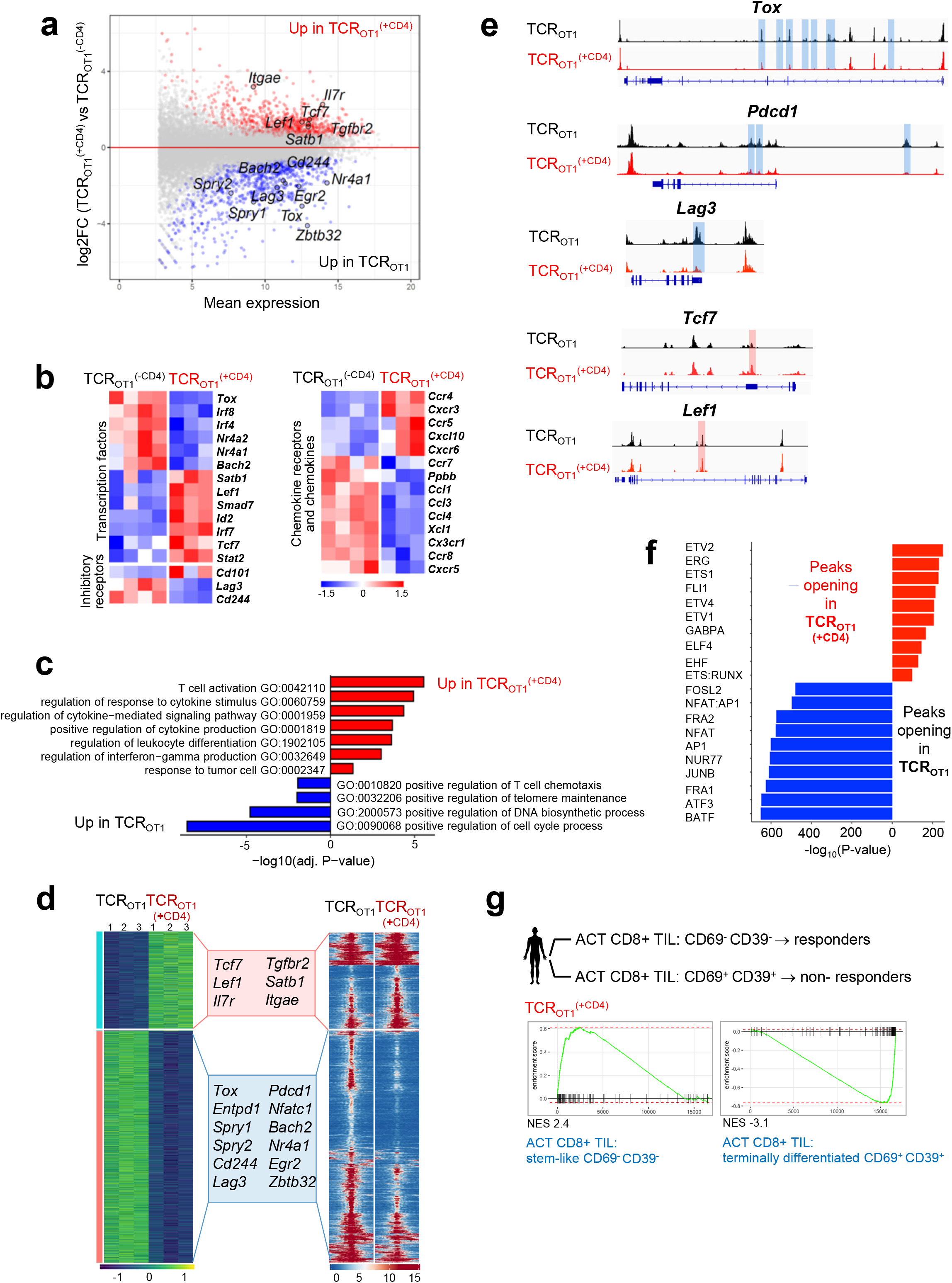
Tumor-specific CD4 T cells transcriptionally and epigenetically reprogram tumor-specific CD8 T cells and prevent terminal differentiation/exhaustion. **a.** MA plot of RNA-seq data showing the relationship between average expression and expression changes of TCR_OT1_ and TCR ^(+CD4)^ TIL. Statistically significantly DEGs (false discovery rate (FDR) < 0.05) are shown in red and blue, with select genes highlighted for reference. **b.** Heat map of RNA-seq expression (normalized counts after variance stabilizing transformation, centered and scaled by row for DEGs) (FDR < 0.05) in TCR_OT1_ and TCR ^(+CD4)^ TIL. **c.** Selected GO terms enriched for genes up-regulated in TCR_OT1_ (blue) and TCR ^(+CD4)^ (red) TIL. **d.** Chromatin accessibility (ATAC-seq); (left) heatmap of log2-transformed normalized read counts transformed with variance stabilization per for regions with differential chromatin accessibility; (right) each row represents one peak (differentially accessible between TCR_OT1_ and TCR ^(+CD4)^ TIL; FDR < 0.05) displayed over a 2-kb window centered on the peak summit; regions were clustered with k-means clustering. Genes associated with the two major clusters are highlighted. **e.** ATAC-seq signal profiles across the *Tox, Pdcd1, Lag3, Tcf7,* and *Lef1* loci. Peaks significantly lost or gained are highlighted in red or blue, respectively. **f.** Top 10 most-significantly enriched transcription factor motifs in peaks with increased accessibility in TCR ^(+CD4)^ TIL (red) or TCR TIL (blue). **g.** Enrichment of gene sets in TCR and TCR_OT1_ (^+CD4^), respectively, described for human tumor infiltrating (TIL) CD8 T cell subsets (CD69-CD39-) stem-like CD8 T cells/TIL (responders) or (CD69+ CD39+) terminally differentiated CD8 T cells/TIL (non-responders) from metastatic melanoma patients receiving *ex vivo* expanded TIL for ACT (S. Krishna *et al*, *Science* 2020). TCR ^(+CD4)^ are enriched in genes observed in CD69-CD39-stem-like T cells/TIL from responders in contrast to TCR_OT1_ which are positively enriched for genes in CD69+ CD39+ terminally differentiated CD8 T cells/TIL from non-responders. NES, normalized enrichment score.

**Figure 3.**
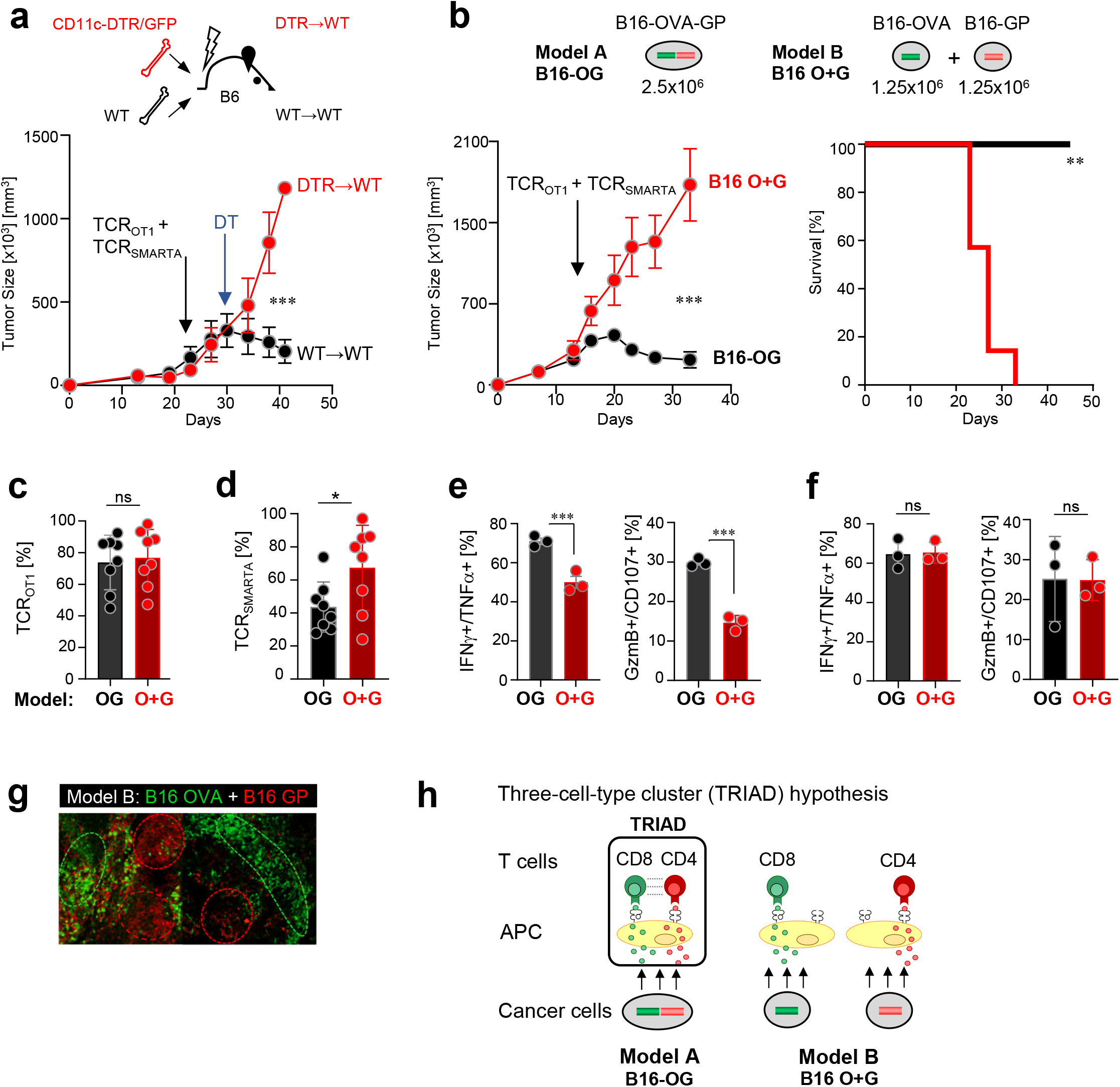
Tumor elimination requires tumor antigen/epitope linkage and unique spatial orientation of tumor-specific CD8 T cells, CD4 T cells and CD11c+ dendritic cells (DC) within tumors. **a.** B16-OG tumor outgrowth in CD11c-DTR/GFP bone marrow (BM) chimeras (scheme, top; DTR→WT or WT→WT) treated with diphtheria toxin (DT). *In vitro* activated TCR_OTI_ and TCR_SMARTA_ were adoptively transferred into lymphodepleted tumor-bearing BM chimeras. 5 days post ACT, mice were treated with DT. Representative of 2 independent experiments (n=3 mice/group). Values are mean ± SEM. Significance is calculated by multiple *t* test. **b.** (Top) Experimental scheme of tumor models A and B: 2.5×10^6^ B16-OG cancer cells (B16 OG; model A) or 1.25×10^6^ B16-OVA (B16-O) mixed with 1.25×10^6^ B16-GP_61-80_ cancer cells (B16 O+G; model B) were transplanted into B6 WT mice. (Bottom), (left) Tumor outgrowth of B16-OG or B16 O+G tumors after TCR_OTI_ and TCR_SMARTA_ ACT. Representative of 2 independent experiments (n=7 mice/cohort). Data are shown as mean ± SEM. Significance is calculated by multiple *t* test. (Right) Kaplan–Meier curve; **p=0.0002; Mantel–Cox test. **c.** Percentage of TCR ^(+CD4)^ (out of total CD8^+^ TIL) 9 days post ACT. **d.** Percentage of TCR_SMARTA_ (out of total CD4^+^ TIL) 9 days post ACT. Data represent 2 pooled, independent experiments (n=8 mice/tumor model). Each symbol represents an individual mouse. **e.** IFNγ, TNFα, CD107, Granzyme B production of TCR ^(+CD4)^ isolated from B16-OG or B16 O+G tumors, or **f.** isolated from tumor-draining lymph nodes of B16-OG or B16 O+G tumor-bearing hosts. Cytotoxic molecules and cytokine production assessed after 4-hr peptide stimulation *ex vivo*. Representative of 2 independent experiments (n=3 mice/tumor). Data are shown as mean ± SEM. *p<0.05, unpaired two-tailed Student’s *t* test. NS, not significant. **g.** Mosaic, clonal growth of B16 OVA-EGFP mixed with B16 GP_61-80_-Cerulean tumor cells (B16 O+G) in B6 WT mice. Shown are confocal microscopy sections of tumors with B16 OVA (green) and B16 GP (red) distinct tumor regions. **h.** Proposed model: Triad formation (three-cell-type clusters; CD8 T cells::CD4 T cells:: APC) form in B16 OG tumors (Model A) where CD8-and CD4-tumor antigens/epitopes are linked and co-presented on the same APC within tumors; tumor-specific CD8 and CD4 T cells engage on same APC; CD4 T cells reprogram CD8 T cells. **Model B:** B16 O+G; triads cannot form due to CD8-and CD4-tumor antigens being presented on distinct APC.

Next, we wanted to investigate *how* TCR_SMARTA_, TCR ^(+CD4)^ and stromal cell interactions cause tumor elimination. To answer this question, we modified our tumor model (**Fig. 3b**): we generated B16 tumor cell lines expressing either the CD8-OVA (B16-O) or CD4-GP (B16-G) tumor antigens. We implanted a mixture of 1.25×10^6^ B16-O and 1.25×10^6^ B16-G cancer cells into WT B6 mice, forming mixed B16 O+G tumors. Control mice received 2.5×10^6^ B16-OG tumor cells as in Figures 1 and 2; thus, both cohorts received the *same* total number (2.5×10^6^) of cancer cells, expressing similar levels of OVA and GP tumor antigens (data not shown). B16 O+G tumors grew with similar kinetics as B16-OG tumors. 2-3 weeks post tumor transplantation, mice received effector TCR_OTI_ and TCR_SMARTA_. 7 days post ACT, equal numbers of TCR_OTI_ and TCR_SMARTA_ TIL were found within progressing B16 O+G and B16-OG tumors (**Fig. 3c, 3d**). Strikingly, despite the same numbers of tumor cells, equal tumor sizes, and same numbers of TCR_OT1_ and TCR_SMARTA_ TIL, mixed B16 O+G tumors continued to grow, in contrast to B16-OG tumors, which ultimately regressed (**Fig. 3b**). TCR_OT1_ TIL isolated from B16 O+G tumors revealed a dysfunctional phenotype similar to those described for TCR_OT1_ transferred without CD4 T cells shown in Figure 1 (**Fig. 3e**). Importantly, these functional differences were only observed within the tumor and not in the tdLN (**Fig. 3f**).

What are the factors and mechanisms that determine tumor progression or regression if numbers of cancer cells and antigen-specific CD8 and CD4 TIL are equal? We hypothesized that a unique spatial organization of cancer cells, CD4 T cells, CD8 T cells, and DC within tumors likely drove CD8 T cell reprogramming and tumor destruction.

### Intratumoral immune triads in mouse and human tumors are required for anti-tumor responses

To define the intratumoral spatial characteristics we conducted confocal microscopic analysis of established B16 O+G tumors. We found regions of either B16-OVA-positive and B16-GP-positive cancer cells, and very few regions that had B16-OVA and B16-GP cancer cells intermingled (**Fig. 3g**). The mosaic-like appearance of distinct tumor regions is a typical feature of clonally growing cancer cells in transplantation tumor models [45]. Consequently, in B16 O+G tumors CD8 or CD4 antigens are largely presented in distinct regions within the tumor and on *distinct* DC/APC (**Model B**), unlike in B16-OG tumors where CD8 and CD4 antigens are co-presented on the *same* DC/APC through epitope linkage (**Model A**) (**Fig. 3h**). Thus, we propose the following model: co-presentation of tumor-specific CD4 and CD8 tumor antigens on the same APC will “force” antigen-specific CD4 and CD8 T cells to form three-cell-type clusters (triads) with APC, and the physical proximity of CD8 T cells with CD4 T cells drives CD4 T cell-mediated CD8 T cell reprogramming and cancer cell destruction (**Model A**). In **Model B**, CD8 and CD4 T cells fail to form triads with APC, CD4 T cells are unable to mediate CD8 T cell reprogramming, ultimately allowing tumors to progress. The concept of a ‘three-cell-type cluster’ was first described in 1987: Mitchison and O’Malley suggested that three-cell-type clusters of CD4 T cell-CD8 T cells-APC were required for the cytolytic response of CD8 T cells in an allogeneic transplant setting [62]. However, little is known about their functional relevance *in vivo* and/or underlying mechanisms.

To determine whether triads are indeed a requisite for tumor elimination, we generated color-coded B16 O+G and B16 O-G tumor models: TCR_SMARTA_ transgenic mice were crossed to EGFP transgenic mice, generating EGFP-expressing TCR_SMARTA_ CD4 T cells; TCR_OT1_ were engineered to express the red fluorescent protein (RFP); CD11c-YFP mice were used as hosts (with yellow fluorescent protein (YFP) under the transcriptional control of the CD11c promoter, thereby YFP-labeling CD11c+ host cells). B16-OG, B16-O, and B16-G cancer cells expressed Cerulean. B16-OG or B16 O+G tumors were established in CD11c-YFP mice and effector TCR_OTI_-RFP^+^ and TCR_SMARTA_-EGFP^+^ adoptively transferred (**Fig. 4a**). Strikingly, 8-9 days post ACT significantly higher numbers of TCR_OT1_::CD11c^+^YFP^+^::TCR_SMARTA_ three-cell-clusters/triads (∼30 interactions/field (or close apposition)) were present in B16-OG tumors, which eventually regressed, in contrast to B16 O+G tumors (∼7 interactions), which eventually progressed (**Fig. 4b**). When normalized to the total number of infiltrating CD11c^+^YFP^+^ cells/field, which remained constant in both tumor models (**Fig. 4c**, right), we observed a 3.5-fold increase of triads in B16-OG tumors (**Fig. 4c**, left). Importantly, dyads, two-cell-interactions between TCR_SMARTA_::CD11c^+^YFP^+^ DC, were not significantly different between B16-OG and B16 O+G (**Fig. 4d**). Thus, CD8 T cell::CD4 T cell::DC triads are associated with tumor-specific CD8 T cell reprogramming and tumor elimination.

**Figure 4.**
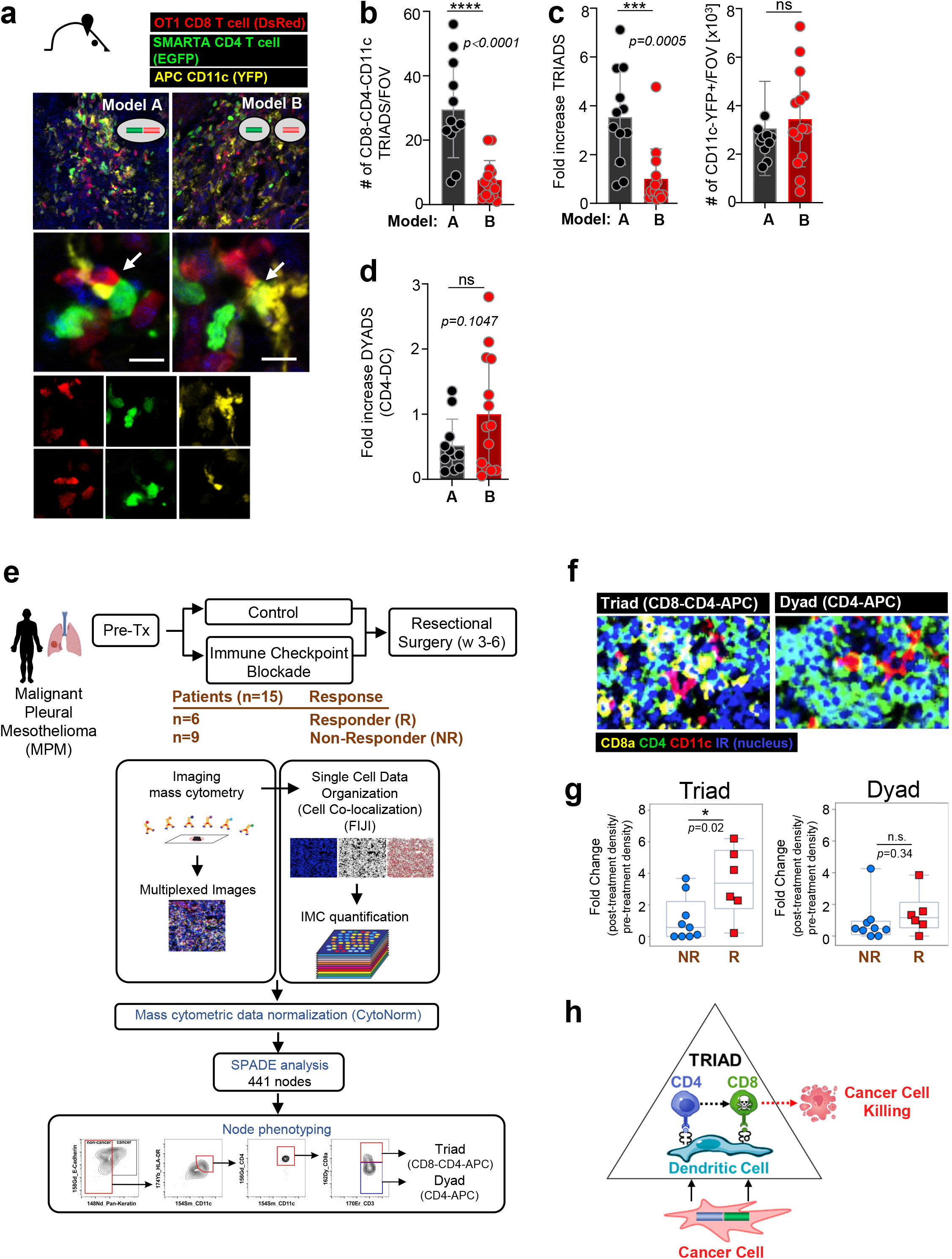
Intratumoral immune triads (three-cell-types clusters; CD8 T cell::CD4 T cell::APC) are required for CD8 T cell reprogramming and tumor elimination. **a.** Color-coded mouse models to determine intratumoral immune triad formation (Models A and B (see Fig. 3)). B16 OG (Model A) or B16 O+G (Model B) tumors were established in CD11c-YFP mice (yellow); effector TCROTI-RFP (red) and TCRSMARTA-EGFP T cells (green) were adoptively transferred into tumor-bearing hosts. Confocal microscopy analysis of frozen tumor tissue sections. Arrows indicate triads. **b.** Numbers of triads per field of view (FOV), and **c.** (left) Fold increase of triads normalized to total numbers of CD11c^+^YFP^+^ cells/FOV (right). c. Quantification of fold increase of numbers of CD4 T cell-DC dyads normalized to total number of infiltrating CD11c^+^YFP^+^ cells/FOV. Each symbol represents an individual frozen tumor section (n=3 mice/group/model). Data are shown as mean ± SEM. *** *P* <0.001, unpaired two-tailed Student’s *t* test. (e.-g). Increased triads in patients with Malignant Pleural Mesothelioma (MPM) treated with checkpoint immunotherapy is associated with pathologic responses. **e.** Treatment regimen and methodology used to determine triads (CD8 T cell::CD4 T cell::APC) and dyads (CD4::APC). Pipeline of co-localization detection by imaging mass cytometry (IMC; see Methods for more details). Briefly, FFPE tumor tissues were stained with 35 target-specific antibodies. Automated cluster detection estimated cluster boundaries by expanding the perimeter of nuclei, identified by Cell ID Intercalator-iridium (191Ir). IMC images were quantified through FIJI, and protein expression data extracted through mean intensity multiparametric measurements performed on individual clusters. Acquired cluster data were normalized with CytoNorm tools, and normalized cytometric data transferred into additional Spanning-tree Progression Analysis of Density-normalized Events (SPADE) to generate automated clustering algorithm and applied cytometric analysis in FlowJo. **f.** Representative multiplexed mass cytometry images of triads and dyads. **g.** Fold change of triads and dyads of pre-and post-immune checkpoint therapy (Tx) density (numbers/mm^2^) in responders (R) and non-responders (NR); *p=0.02; n.s. p=0.34 (not significant). **h.** Proposed model of TRIAD-associated cancer elimination.

Next, we asked whether CD8 T cell::CD4 T cell::APC triads could be associated with clinical responsiveness in humans. As clinical data assessing spatial characteristics of immune cells within tumors of ACT-treated patients was not available, we turned to patients treated with immune checkpoint blockade (ICB) therapy; ICB therapies have shown efficacy in some cancer patients and cancer types, however most patients remain refractory. The underlying mechanisms determining ICB resistance or responsiveness, as well as predictive biomarkers, remain poorly defined. We assessed the spatial orientation of CD8 T cells, CD4 T cells and APC in patients with malignant pleural mesothelioma (MPM) undergoing ICB therapy [63]. Patients were randomized and treated with Durvalumab (anit-PDL1) mono-or Durvalumab and Tremelimumab (anti-CTLA4) combination therapy. A no ICB group was included as a control cohort. Tumor tissues were obtained both before and after ICB treatment [63]. Evaluable tumors, before and after ICB were available for 15 patients receiving ICB. Out of the 15 patients, 6 patients showed a pathologic response (R; Responders) while 9 patients did not (NR; Non-Responders) (**Fig. 4e**). Imaging mass cytometry (IMC) and time-of-flight mass cytometry (CyTOF) were performed on all 15 patients’ pre-and post-treatment tumor tissues using 35 markers to determine co-localization of non-T_REG_ CD4 T cells, CD8 T cells, and CD11c+ APC, including the presence of dyads (CD4::APC or CD8::APC) and triads (CD4::CD8::APC) (**Fig. 4e** and **4f**). Strikingly, while neither numbers of tumor-infiltrating CD8 T cells, nor CD4::APC or CD8::APC dyads were associated with a pathologic response and ICB responsiveness, triads were able to demarcate responders from non-responders (**Fig. 4g**). Our studies reveal triads as critical determinants for anti-tumor immunity and ICB responsiveness in patients with MPM.

## DISCUSSION

Here, we demonstrate a new role for CD4 T cells during the effector phase of cytotoxic CD8 T cell-elimination of solid tumors in the setting of ACT. CD4 T cell reprogramming of CD8 T cells and cancer cell elimination is strictly dependent on the formation of immune triads, tumor-specific CD8 T cells and CD4 T cells co-engaged with the same DC, and not on CD4 T cell engagement with cancer cells, important given that most epithelial cancers do not express MHC class II. We demonstrate that the spatial positioning of CD8 and CD4 T cells within tumors, and not the number of intratumoral tumor-specific CD8 and CD4 T cells, is the critical determinant of effective anti-tumor immunity and ACT efficacy. Our data may provide clues as to why ACT clinical trials utilizing predominantly tumor-reactive CD8 T cells have shown only limited responses for the treatment of solid tumors.

It is well established that CD4 T cells are required for CD8 T cell effector differentiation. However, studies have mainly focused on CD4 T cell ‘help’ of naïve CD8 T cells during the priming/activation phase and memory formation in infection and vaccination settings [31, 42, 64-66]. The importance of CD8-CD4 T cell co-operation during the priming/activation phase was elegantly described by the Germain group, demonstrating that nonrandom, chemokine-driven (CCL3, CCL4) recruitment of CCR5+ naïve, antigen-specific CD8 T cells to sites of antigen-specific DC-CD4 T cell interactions within antigen-draining lymph nodes led to optimal CD8 T cell responses during vaccination and early infections [30]. CD4 T cells license DC through CD40L-CD40 interactions, enhancing B7 and CD70 expression on DC; CD28-and CD27-expressing antigen-specific CD8 T cells (ligands for B7 and CD70, respectively) receive optimal co-stimulatory signals when engaging with DC-CD4 T cells and/or abundant IL2 produced by CD4 T cells. Vaccines relying only on short, single MHC class I-restricted peptides showed reduced clinical benefits compared to synthetic long peptide vaccine platforms containing both MHC class I and class II epitopes, highlighting the importance of guided CD8 and CD4 cooperation [42-46]. Here, we discover that CD4 T cells and triads are critical for cancer cell elimination by cytolytic effector CD8 T cells: antigen-specific CD4 T cells within tumors reprogram antigen-specific effector CD8 T cells, repressing terminal differentiation and preserving stem-like features and effector function. Physical proximity of CD8 T cells with CD4 T cells likely enforces chemokine and/or cytokine signaling, or direct receptor-ligand interactions needed for CD8 T cell reprogramming. Interestingly, chemokine receptors such as *Ccr5*, *Ccr4* and *Ccr2* were upregulated on TCR_OT1(+CD4)_ encountering DC-CD4 T cells, as well as *Il2rg* and *Ifngr1*. Future studies must determine the precise mechanisms by which CD8 T cells resist T cell exhaustion and mediate cancer destruction. Our finding that triads (but not dyads) were associated with a pathogenic anti-tumor response in ICB-treated patients with malignant pleural mesothelioma, suggests that intratumoral immune triads may also be critical for anti-tumor responses in non-ACT settings. Interestingly, and congruent with our findings, a recent study demonstrated that dendritic cell–CD4 T helper cell niches enable CD8 T cell differentiation in patients with hepatocellular carcinoma following PD-1 blockade [67].

Our study reveals a previously unappreciated role of unique cell-cell interactions and spatial positioning within tumors where tumor-specific CD4 T cells empower tumor-specific CD8 T cells to eliminate solid tumors in adoptive T cell therapy. MHC class II-restricted neoantigens or self/tumor antigens and tumor-specific CD4 T cells have been described in human cancers [48-50]. Designing therapeutic interventions that enforce the formation of CD4-CD8-DC triads in tumors might be powerful strategies for the treatment of cancers, including for ICB-, vaccine-and ACT-approaches.

## MATERIALS AND METHODS

### Mice

B6 mice (C57BL/6J), TCROTI (C57BL/6-Tg(TcraTcrb)1100Mjb/J), TCRSMARTA (B6.Cg-Ptprca Pepcb Tg(TcrLCMV)1Aox/PpmJ), CD11c-YFP (B6.Cg-Tg(Itgax-Venus)1Mnz/J), CD11c-DTR-GFP (B6.FVB-1700016L21RikTg(Itgax-DTR/EGFP)57Lan/J), GFP transgenic (C57BL/6-Tg(CAG-EGFP)1Osb/J), B6 Thy1.1 (B6.PL-Thy1a/CyJ), and B6 CD45.1 (B6.SJL-Ptprca Pepcb/BoyJ) mice were purchased from the Jackson Laboratory. TCRSMARTA mice were crossed to Thy.1.1 mice to generate TCRSMARTA Thy1.1 mice; for Figure 4 imaging studies, TCRSMARTA Thy1.1 mice were crossed to GFP-transgenic mice to generate TCRSMARTA Thy1.1 GFP mice. TCROTI (Thy1.2) mice were crossed to CD45.1 mice to generate TCROTI CD45.1 mice. Both female and male mice were used for experimental studies. Donor and host mice were age-and sex-matched; mice were 7-12 weeks old. All mice were bred and maintained in the animal facility at Memorial Sloan Kettering Cancer Center (MSKCC). Experiments were performed in compliance with the MSKCC Institutional Animal Care and Use Committee (IACUC) regulations.

### Antibodies and Reagents

Fluorochrome-conjugated antibodies were purchased from BD Biosciences, eBioscience, and Biolegend. The OVA257-264 and GP61–80 peptides were purchased from GenScript.

### Intracellular cytokine staining

Intracellular cytokine staining was performed using the Foxp3 staining kit (BD Biosciences) following the manufacturer’s protocol. Briefly, T cells isolated from lymph nodes or tumors were mixed with 3×10^6^ congenically marked B6 splenocytes and incubated with 1 μg/mL of OVA peptide and/or 2 μg/mL of GP peptide for 4-5h at 37°C in the presence of GolgiPlug (BD Biosciences). After staining for cell surface molecules, cells were fixed, permeabilized and stained with antibodies against IFNγ (XMG1.2) and TNFα (MP6-XT22).

### Flow Cytometric Analysis

Flow cytometric analysis was performed using Fortessa X20. Cells were sorted using BD FACS Aria (BD Biosciences) at the MSKCC Flow Core Facility. Flow data were analyzed with FlowJo v.10 software (Tree Star Inc.).

### Generation of plasmids and tumor cell lines

#### Tumor antigen-encoding pMFG-Cerulean vectors

pMFG-OVA257-264-Cerulean, pMFG-GP61–80-Cerulean, and pMFG-OVA257-264-GP61–80-Ceruelan plasmids were constructed by inserting annealed oligonucleotides encoding triple SIINFEKL-AAY repeats, GLKGPDIYKGVYQFKSVEFD, or (SIINFEKL-AAY)3-P2A-GLKGPDIYKGVYQFKSVEFD, respectively, into the NcoI-linearized pMFG-Cerulean vector, as previously described [45]. Restriction enzymes were purchased from New England Biolabs. All constructs were verified by sequence analysis. Phoenix packaging cells (ATCC) were transfected with pMFG constructs; supernatants were used to transduce B16-F10 mouse melanoma tumor cell line to generate B16-F10 OVA257-264-Cerulean, B16-F10-GP61-80-Cerulean and B16-F10 OVA257-264-GP61-80-Cerulean, respectively [45]. Transduced bulk cell lines were sorted for similar Cerulean expression levels.

### *In vitro* T cell activation

For the generation of effector TCROT1 CD8 T cells and TCRSMARTA CD4 T cells, single-cell suspensions were prepared from spleens of TCROT1 and TCRSMARTA transgenic mice and cultured *in vitro* in RPMI 1640 medium supplemented with 10% FBS, 100 IU/ml penicillin, 100 mg/ml streptomycin, nonessential amino acids, 1 mM sodium pyruvate, and 20 mM HEPES, together with 1 μg/mL of OVA257-264 peptide or 2 μg/mL of GP61–80 peptide, respectively, at a concentration of 4-5×10^6^ splenocytes/ml in the presence of 50U/mL IL-2 for 4 days.

### Adoptive T cell transfer

For adoptive transfer studies, 2.5×10^5^ *in vitro* activated TCROT1 (CD45.1) and/or 5×10^5^ *in vitro* activated TCRSMARTA (Thy1.1) were transferred (i.v.) into tumor-bearing WT B6 mice at indicated time points post tumor transplantation (approximately 2-3 weeks post tumor implantation). Tumor-bearing mice were treated with cyclophosphamide (180mg/kg), and 24h later *in vitro* activated TCROT1 CD8 T cells and/or TCRSMARTA CD4 T cells were adoptively transferred. At indicated time points, adoptively transferred T cells were isolated from tumor-draining lymph nodes and tumors and prepared for downstream analyses.

### B16 and MCA 205 transplantation tumor models

2.5×10^6^ B16 OVA257-264-GP61-80 (B16 OG) tumor cells, or a mixture of 1.25×10^6^ B16 OVA257-264 (B16 O) + 1.25×10^6^ B16 GP61-80 (B16 G) tumor cells (B16 O+G), or MCA OVA257-264-GP61-80 tumor cells were injected subcutaneously into mice. Antigen-specific T cells were adoptively transferred into tumor-bearing mice as described in text and figure legends. For outgrowth experiments, tumors were measured manually with a caliper. Tumor volume was estimated with the formula (L x W x H)/2.

### Generation of bone marrow chimeras and depletion of dendritic cells *in vivo*

B6 WT (CD45.1) mice were irradiated twice with 600 cGy, 6 hours apart. 12-18 hours later, bone marrow (BM) was isolated from femurs and tibias of CD11c-DTR/GFP (CD45.2) mice, and 5–8×10^6^ BM cells were injected i.v. into irradiated CD45.1 mice. BM chimeric were given antibiotics (trimethoprim-sulfamethoxazole) for 2 weeks. BM chimeric were analyzed for successful engraftment and BM reconstitution 6-8 weeks later. For conditional DC depletion, CD11c-DTR/GFP BM chimeric mice were injected (i.p.) with 4–5 ng/g body weight diphtheria toxin (DT, Sigma-Aldrich) every other day for 14 days.

### Generation of B16 -*I-A^b^*-deficient tumor cell line

The B16 tumor cells were subjected to CRISPR/Cas9-mediated knockout of *I-A^b^* by transient transfection of a plasmid encoding both Cas9 nuclease and single guide (sg) RNA targeting the *I-A^b^*locus, as well as GFP reporter gene. 2.5×10^5^ B16 cells were plated and transfected with 2μg of Cas9-and sgRNA-encoding plasmid DNA using Lipofectamine 3,000 (Invitrogen) following the manufacturer’s protocol. 3 days post transduction, GFP+ cells were FACS-sorted. Deletion of *I-A^b^*was confirmed by treating GFP+ B16 *I-A^b^* cells with 20 U/ml IFNγ for 48h, followed by flow cytometric analysis of I-A^b^ expression.

### Color-coded tumor model and adoptive transfer of color-coded T cells

CD11c-YFP transgenic mice were injected subcutaneously with 2.5×10^6^ (B16 OG) tumor cells or a mixture of 1.25×10^6^ B16-O + 1.25×10^6^ B16-G tumor cells (B16 O+G). To generate color coded TCROT1 CD8 T cells, TCROT1 splenocytes were transduced to express tRFP using retroviral transduction as previously described [68]. Briefly, Platinum-E cells (ATCC) were transfected with a tRFP-encoding retroviral vector using the Mirus TransIT-LT1 reagent (catalog no. 2305). Viral supernatant was supplemented with polybrene and added to TCROT1 splenocytes, and the cells were transduced via spinfection on two consecutive days. To generate color-coded TCRSMARTA CD4 T cells, splenocytes from TCRSMARTA GFP transgenic mice were used and activated as described above. Tumor-bearing mice were treated with cyclophosphamide (180mg/kg) one day before ACT, and *in vitro* activated 2.5+10^5^ TCROT1 tRFP+ CD8 T cells and 4X10^5^ cells TCRSMARTA EGFP CD4 T cells were transferred (i.v.) into tumor-bearing mice.

### Immunofluorescence staining and confocal imaging

For confocal microscopy analysis, pieces of established tumors were excised and fixed for 18-24 hours in 4% paraformaldehyde solution, followed by dehydration in 20% sucrose, and then embedded in OCT, and stored at −80°C. 30-μm-thick frozen sections were cut on a CM3050S cryostat (Leica) and adhered to Superfrost Plus slides (Thermo Fisher Scientific). Nuclei were labeled using DAPI (Sigma). Slides were mounted with ProLong Diamond Antifade Mountant (Invitrogen) and analyzed on a Leica TCS SP8 confocal microscope. Fiji Is Just ImageJ (FIJI) was utilized for image analysis. 3D reconstitution was performed, and triple contacts/triads were assessed based on color-coded immune subset identification. Analyses was performed as a blinded outcome assessment. To quantify double contacts, after thresholding and binarization of images, the function “analyze particles” has been applied. To precisely estimate only events showing double contact, the mathematical function “AND” was used.

### Isolation of adoptively transferred T cells from downstream analyses

Lymph nodes were mechanically disrupted with the back of a 3-mL syringe, filtered through a 100-μm strainer, and red blood cells (RBC) were lysed with ammonium chloride potassium buffer. Cells were washed twice with cold RPMI 1640 media supplemented with 2μM glutamine, 100U/mL penicillin/streptomycin, and 3% fetal bovine serum (FBS). Tumor tissue was mechanically disrupted with a glass pestle and a 150-μm metal mesh in 5mL of cold HBSS with 3% FBS. Cell suspension was filtered through 70-μm strainers. Tumor homogenate was spun down at 400*g* for 5 minutes at 4°C. Pellet was resuspended in 15 mL HBSS with 3% FBS, 500 μl (500U) heparin, and 8 mL isotonic Percoll (GE), mixed by several inversions, and spun at 500*g* for 10 min at 4°C. Pellet was lysed with ammonium chloride potassium buffer and cells were further processed for downstream applications.

### Sample Preparation for RNA-Seq and ATAC-Seq

TCROT1 CD8 T cells were isolated from tumors (see above); cells were stained for CD8α (clone 53-6.7, eBioscience) and CD45.1^+^(clone A20, Biolegend). CD8^+^CD45.1^+^ cells were sorted by FACS. For RNA-seq, T cells were directly sorted into Trizol LS reagent (Invitrogen, catalog no. 10296010) and stored at - 80℃. For ATAC-seq, sorted T cells were resuspended in cold FBS with 10% DMSO and stored at -80℃.

### RNA-seq

RNA from sorted cells was extracted using the RNeasy Mini Kit (Qiagen; catalog no. 74104) according to instructions provided by the manufacturer. After RiboGreen quantification and quality control by an Agilent BioAnalyzer, total RNA underwent amplification using the SMART-Seq v4 Ultra Low Input RNA Kit (Clontech), and amplified cDNA was used to prepare libraries with the KAPA Hyper Prep Kit (Kapa Biosystems). Samples were barcoded and run on a HiSeq 2500 in a 50-bp/50-bp paired-end run with the HiSeq SBS Kit v4 (Illumina). An average of 50 million paired reads were generated per sample.

### ATAC-seq

Profiling of chromatin accessibility was performed by ATAC-seq as previously described (Buenrostro et al., 2013). Briefly, viably frozen, sorted T cells were washed in cold PBS and lysed. The transposition reaction was incubated at 42°C for 45 min. The DNA was cleaned with the MinElute PCR Purification Kit (Qiagen; catalog no. 28004), and material was amplified for five cycles. After evaluation by real-time PCR, 7–13 additional PCR cycles were done. The final product was cleaned by AMPure XP beads (Beckman Coulter, catalog no. A63882) at a 1× ratio, and size selection was performed at a 0.5× ratio. Libraries were sequenced on a HiSeq 2500 or HiSeq 4000 in a 50-bp/50-bp paired-end run using the TruSeq SBS Kit v4, HiSeq Rapid SBS Kit v2, or HiSeq 3000/4000 SBS Kit (Illumina). An average of 100 million paired reads were generated per sample.

### Bioinformatics methods

The quality of the sequenced reads was assessed with FastQC and QoRTs (for RNA-seq samples; Hartley and Mullikin, 2015; Andrews, 2010). Unless stated otherwise, plots involving high-throughput sequencing data were created using R version 4.1.0 (R Core Team, 2017) and ggplot2 (Wickham, 2016).

### RNA-seq data

DNA sequencing reads were aligned with default parameters to the mouse reference genome (GRCm38.p6) using STAR v2.6.0c (Dobin et al., 2013). Gene expression estimates were obtained with featureCounts v1.6.2 using composite gene models (union of the exons of all transcript isoforms per gene) from Gencode (version M17; Liao et al., 2014).

### DEGs

DEGs were determined using DESeq2 v1.34.0 with Wald tests with a q-value cutoff of 0.05 (Benjamini– Hochberg correction).

### Heatmaps

Heatmaps in Fig. 2b were created using DESeq2 normalized read counts after variance stabilizing transformation of genes identified as differentially expressed by DESeq2. Rows were centered and scaled.

### Pathway and GO term enrichment analyses

Gene set enrichment analyses (Fig. 2g and Suppl. Fig 1) were done using fgsea v1.20.0 [69] with the fgseaMultilevel function. Genes were ranked based on the DESeq2 Wald statistic. Gene sets with an FDR < 0.05 were considered enriched.

Gene ontology analysis was performed on up-and down-regulated DEGs using the clusterProfiler v4.2.2 R package [70]. Only GO categories enriched using a 0.05 false discovery rate cutoff were considered.

### ATAC-seq data

#### Alignment and creation of peak atlas

Reads were aligned to the mouse reference genome (version GRCm38) with BWA-backtrack v0.7.17 (Li and Durbin, 2009). Post-alignment filtering was done with samtools v1.8 and Picard tools v2.18.9 (Li et al., 2009) to remove unmapped reads, improperly paired reads, nonunique reads, and duplicates. Peaks were called with MACS2 v2.1.1 (Liu, 2014), and peaks with adjusted P values smaller than 0.01 were excluded.

Consensus peak sets were generated for each condition if a peak was found in at least two replicates. Reproducible peaks from each condition were merged with DiffBind v3.4.11 to create an atlas of accessible peaks, which was used for downstream analyses. The peak atlas was annotated using the ChIPseeker v1.30.3 [71] and TxDb.Mmusculus.UCSC.mm10.knownGene [Bioconductor Core Team and Bioconductor Package Maintainer (2019). TxDb.Mmusculus.UCSC.mm10.knownGene: Annotation package for TxDb object(s). R package version 3.10.0.]. Blacklisted regions were excluded (https://sites.google.com/site/anshulkundaje/projects/blacklists).

### Differentially accessible regions

Regions where the chromatin accessibility changed between different conditions were identified with DESeq2 v1.34.0, and only Benjamini–Hochberg corrected P values < 0.05 were considered statistically significant.

### Coverage files

Genome coverage files were normalized for differences in sequencing depth (RPGC normalization) with bamCoverage from deepTools v3.1.0. Replicates were averaged together using UCSC-tools bigWigMerge. Merged coverage files were used for display in Integrated Genomics Viewer shown in Fig. 2e.

### Heatmaps

Heatmaps based on the differentially accessible peaks identified between TCROT1 and TCR ^(+CD4)^ as shown in Fig. 2d were created using profileplyr v1.10.2 (T. Carroll and D. Barrows (2021). profileplyr: Visualization and annotation of read signal over genomic ranges with profileplyr. R package version 1.10.2.) and ComplexHeatmap v2.15.1 [72], by binning the region +/− 1kb around the peak summits in 20bp bins. To improve visibility, bins with read counts greater than the 75th percentile + 1.5*IQR were capped at that value.

### Motif analyses

For identifying motifs enriched in differentially accessible peaks, we utilized HOMER via marge v0.0.4 ([73]; and [Robert A. Amezquita (2021). marge: API for HOMER in R for Genomic Analysis using Tidy Conventions. R package version 0.0.4.9999]). HOMER was run separately on hyper-or hypo-accessible peaks with the flags -size given and -mask. Motifs enriched in hyper-or hypo-accessible peaks were determined by comparing the rank differences (based on P value). The consensus peakset identified by DiffBind was used as the background set.

### Human Data (Fig. 4e-4g)

#### Trial, Patients, Study Design

For more details on patients, study design, and methodology see Hyun-Sung Lee *et al* [63]. Briefly, this was a phase II, prospective, randomized window-of-opportunity trial completed at Baylor College of Medicine that enrolled patients with surgically resectable MPM (NCT02592551). Eligible patients underwent a staging procedure that included cervical mediastinoscopy with mediastinal lymph node biopsies and diagnostic laparoscopy with peritoneal lavage and peritoneal biopsies. Thoracoscopy with tumor biopsies was performed for the purpose of the trial. Patients without pathologic nodal or peritoneal disease were randomly assigned in a 2:2:1 ratio to receive (i) one dose of durvalumab (10 mg/kg i.v.), (ii) one dose of durvalumab (1,500 mg) plus one dose of tremelimumab (75 mg i.v.), or (iii) no ICB. ICB was administered 3 days to 3 weeks following the staging procedure and surgical resection was performed 3 to 6 weeks after ICB by extended pleurectomy/decortication (P/D) or extrapleural pneumonectomy (EPP). Tumor and blood were obtained before and after ICB (at thoracoscopy and resection, respectively).

#### Methods

Cancer specimens were processed into single-cell suspensions, fresh frozen tissue preparations, samples cryopreserved in optimal cutting temperature (OCT) compound, and formaldehyde-fixed paraffin-embedded tissues (FFPE).

##### Imaging mass cytometry (IMC)

FFPE tissue samples were sectioned at a 5-μm thickness for IMC. FFPE tissues on charged slides were stained with 1:100 diluted antibody cocktails (concentration of each antibody=0.5mg/mL) as recommended by the user’s manual. The slides were scanned in the Hyperion Imaging System (Fluidigm). They were scanned at least four regions of interest in >1mm2 at 200 Hz.

##### SIMC analysis

Fiji was used for cell segmentation and conversion of imaging data into flow cytometric data, with the advantage of fast, robust, unsupervised, automated cell segmentation method. 32-bit TIFF stacked images were loaded in Fiji and novel method of automated cell segmentation that estimates cell boundaries by expanding the perimeter of their nuclei, identified by Cell ID intercalator iridium (191Ir) was used as described in more detail in Hyun-Sung Lee *et al* [63]. Once images from the IMC methodology were acquired, images were quantified through FIJI’s threshold and watershed tools. Protein expression data were then extracted at the single-cell level through mean intensity multiparametric measurements performed on individual 10 cells and acquired single-cell data were transferred into additional cytometric analysis in FlowJo V10 software (FlowJo, LLC, OR). All protein markers in quantified IMC data are adjusted with 191Ir and 193Ir nucleus intensities and normalized with CytoNorm across IMC regions of interests, a normalization method for cytometry data applicable to large clinical studies that is plugged-in FlowJo. CytoNorm allows reducing mass cytometry signal variability across multiple batches of barcoded samples. Normalized IMC data are combined by using FlowJo. For <u>CyTOF,</u> please see Hyun-Sung Lee *et al* [63].

### Statistical analyses

Statistical analyses on flow cytometric data were performed using unpaired two-tailed Student’s *t* tests (Prism 7.0, GraphPad Software). A *P* value of < 0.05 was considered statistically significant. All other statistical testing methods are described in figure legends.

## Acknowledgements

We thank the members of the Schietinger lab for helpful discussions, and M. Philip (Vanderbilt University) for critical reading of the manuscript. This work was supported by NIH grants DP2CA225212 and R01CA269733 (A.S.), Lloyd Old STAR Award of the Cancer Research Institute (A.S.), AACR-Bristol Myers Squibb Midcareer Female Investigator Award (A.S.), Pershing Square Sohn Award (A.S.), Josie Robertson Young Investigator Award (A.S.), the Weill Cornell Medicine Core Laboratories Center (P.Z., D.B), Ludwig Cancer Center Postdoctoral Fellowship (G.E.C.); NIH R37 MERIT Award R37CA248478 (B.M.B), Cancer Prevention and Research Institute of Texas Grant CPRIT RP200443 (H.S.L.), Department of Defense Peer Reviewed Cancer Impact Award CA210522 (H.S.L.).We acknowledge the use of the Integrated Genomics Operation Core, funded by the NCI Cancer Center Support Grant (P30 CA08748), Cycle for Survival, and the Marie-Josée and Henry R. Kravis Center for Molecular Oncology.

## Author Contributions

G.E.C. and A.S. conceived and designed the study; G.E.C, carried out experiments, analyzed and interpreted data. A.D. assisted with mouse breeding. P.Z. and D.B. performed computational analyses. A. Scrivo conduced microscopy analyses. For human study: conceptualization (B.M.B., H.S.L. and M.H.). H.S.L. and B.M.B: data curation, analyses, visualization, methodology. G.E.C. and A.S. wrote the manuscript, with all authors contributing to the writing and providing feedback.

---

**Supplementary Figure 1:**
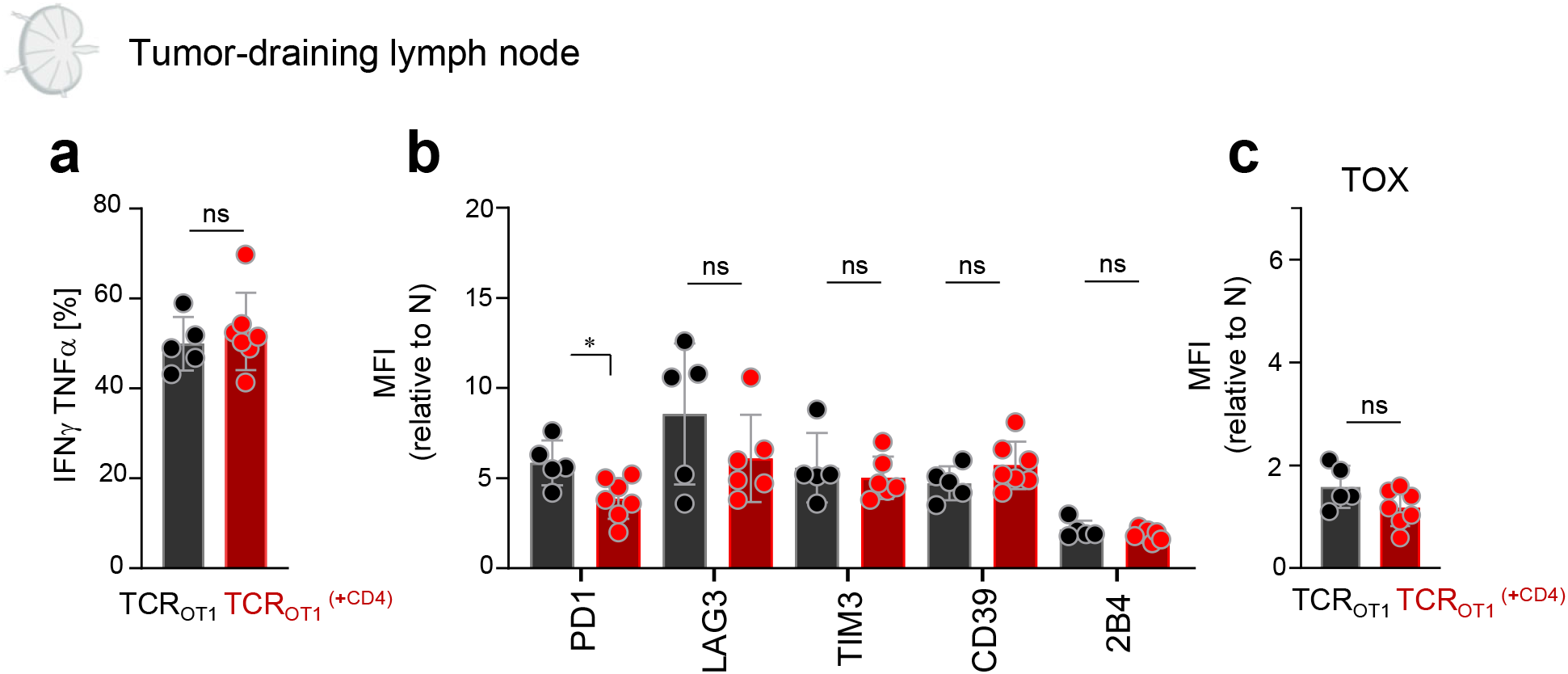
**a.** IFNγ and TNFα production, **b.** inhibitory receptor expression, and **c.** TOX expression of TCR_OTI_ isolated from tumor-draining lymph nodes of B16-OG tumor-bearing mice 8-9 days post transfer +/-TCR_SMARTA_. Cytokine production was assessed after 4-hr peptide stimulation *ex vivo*. Data show 2 pooled independent experiments (n=5-7). Data are shown as mean ± SEM. *p<0.05, using unpaired two-tailed Student’s *t* test. ns, not significant.

**Supplementary Figure 2:**
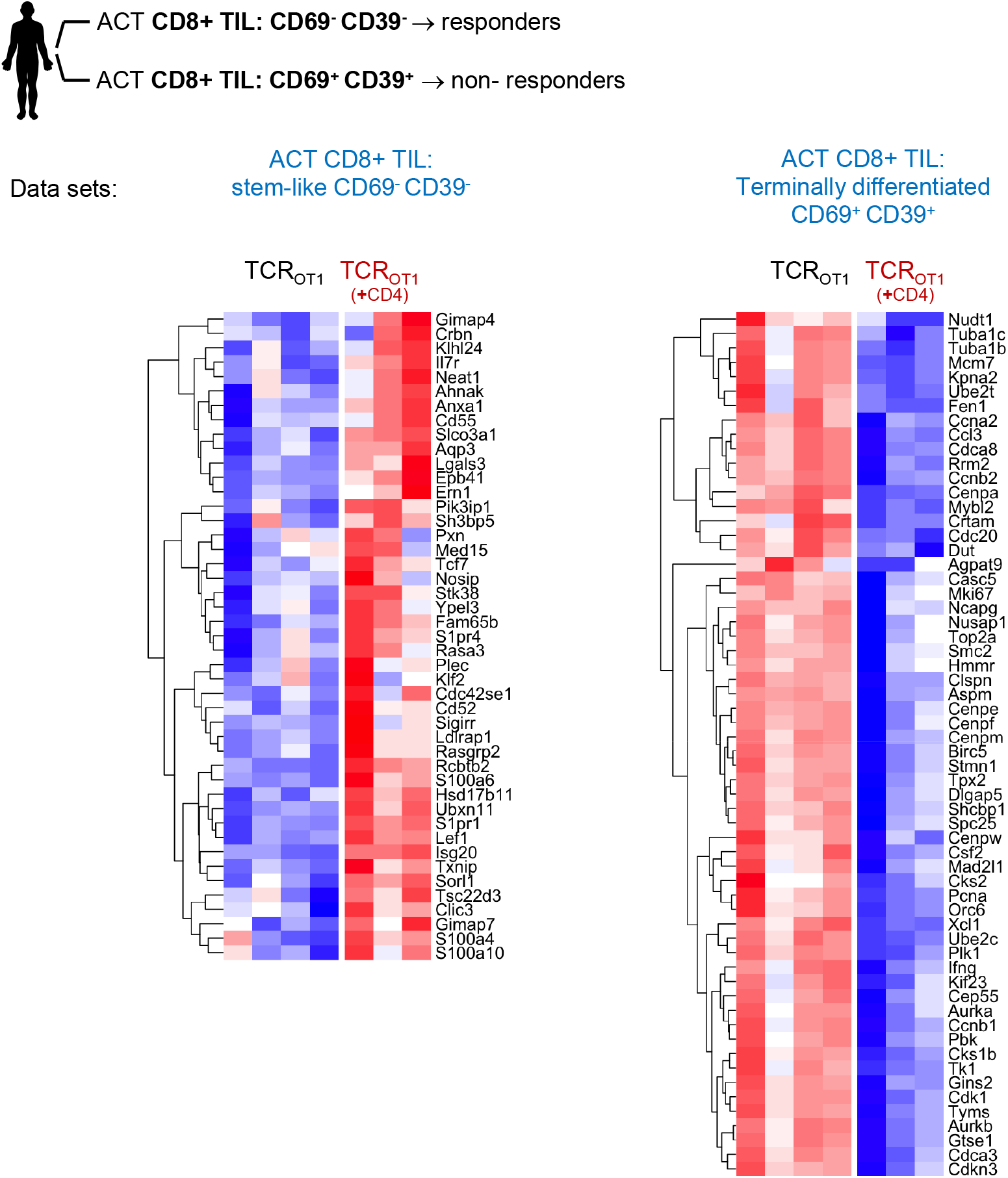
Enrichment of gene sets in TCR_OT1_ and TCR_OT1_ (+CD4), respectively, described for human tumor infiltrating (TIL) CD8 T cells from metastatic melanoma patients receiving *ex vivo* expanded CD8+ TIL in in adoptive T cell transfers (ACT) (S. Krishna *et al*, *Science* 2020). ACT responders contained CD69-CD39-stem-like CD8+ TIL, which were lacking in ACT-non-responders. ACT non-responders contained CD69+ CD39+ terminally differentiated CD8+ TIL. TCR_OT1_ (+CD4) are enriched in genes observed in CD69-CD39-stem-like T cells/TIL and are negatively enriched for genes from CD69+ CD39+ terminally differentiated CD8 T cells/TIL. Significantly differentially expressed, enriched genes are shown. See also main Figure 2g.

**Supplementary Figure 3:**
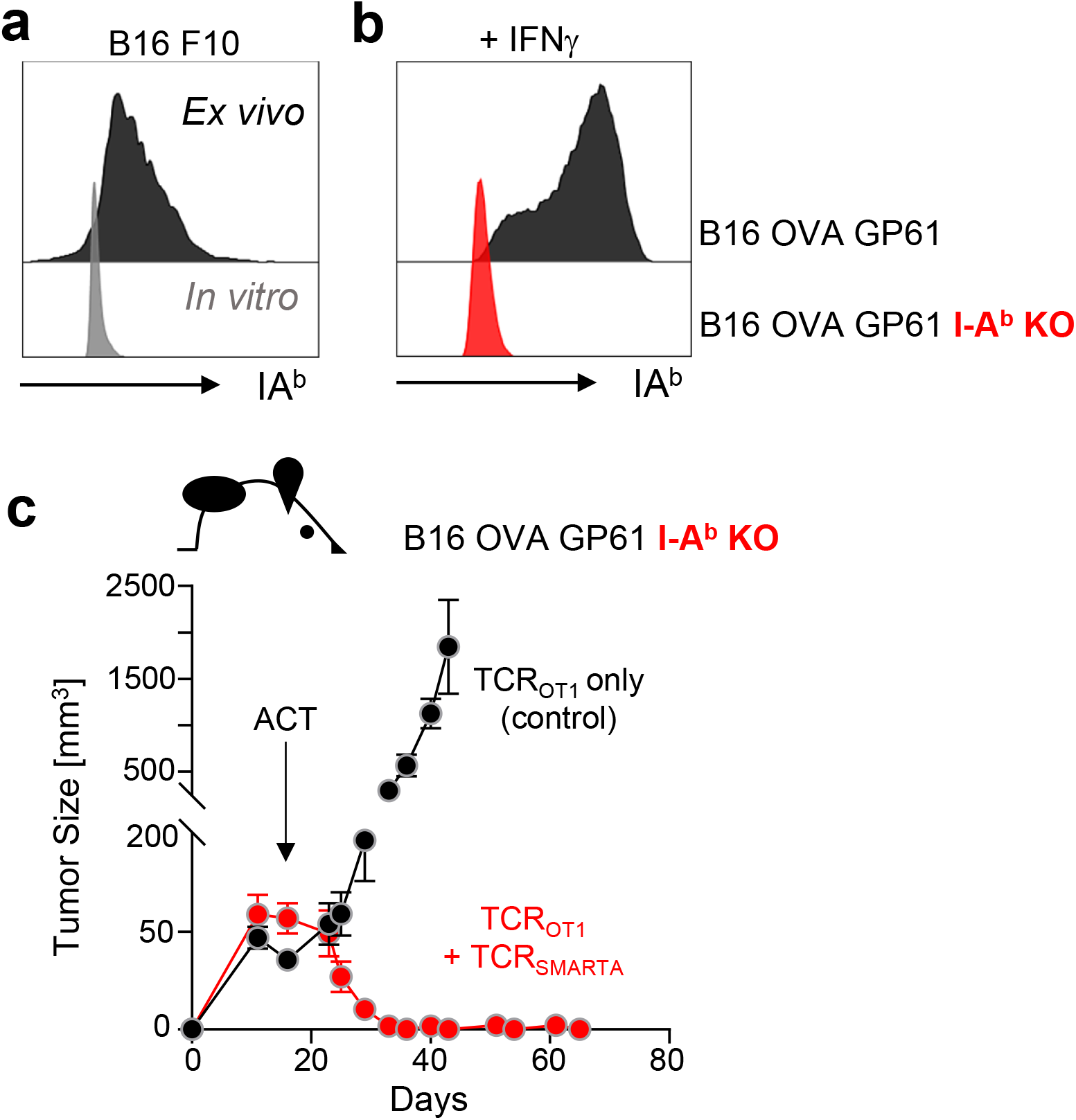
**a.** Flow cytometric analysis of MHC class II I-A^b^ expression on parental B16 tumor cells cultured *in vitro* (grey) or after isolation from tumor bearing B6 WT mice *ex vivo* (black). **b.** I-A^b^ expression on B16-OG tumor cells (parental; black) or CRISPR/Cas9 gene-edited B16 OG *I-A^b^*-deficient cells (KO; red) after 48 hours IFNγ treatment *in vitro*. **c.** Outgrowth of B16-OG *I-A^b^*-deficient tumors in B6 WT mice receiving adoptively transferred *in vitro* activated TCR_OT1_ and TCR_SMARTA_ (red) or TCR_OT1_ only (black).

## Notes

### Competing Interest Statement

The authors have declared no competing interest.

